# Combinations comprising dual β-lactams and a β-lactamase inhibitor achieve optimal synergistic inhibition of *Mycobacterium abscessus* growth

**DOI:** 10.1101/2025.01.20.633926

**Authors:** Binayak Rimal, Yi Xie, Chandra M. Panthi, Kaylyn L. Devlin, Kimberly E. Beatty, Gyanu Lamichhane

**Affiliations:** Division of Infectious Diseases, Department of Medicine, School of Medicine, Johns Hopkins University, Baltimore, MD 21231, USA; Department of Chemical Physiology and Biochemistry, Oregon Health Sciences University, Portland, OR 97239, USA; Center for Nontuberculous Mycobacteria and Bronchiectasis, School of Medicine, Johns Hopkins University, Baltimore, MD 21231, USA

**Keywords:** *Mycobacterium abscessus*, β-lactams, β-lactamase, β-lactamase inhibitor, dual β-lactam, dual β-lactamase inhibitor

## Abstract

The historical model, which posits that β-lactams inhibit bacterial growth while β-lactamase inhibitors (BLIs) merely protect β-lactams from enzymatic degradation, fails to fully explain their activity against *Mycobacterium abscessus* (*Mab*). This study demonstrates that synergistic effects extend beyond the traditional one β-lactam+one BLI paradigm, refuting the oversimplified mechanistic framework. First, β-lactam-based BLIs such as clavulanic acid, sulbactam, and tazobactam exhibit intrinsic antibacterial activity against *Mab*. These agents synergized not only with β-lactams but also with one another, undermining their historical classification as mere β-lactamase inhibitors. The data indicate that their activity is not limited to inhibiting β-lactamases but extends to directly targeting critical bacterial processes. Second, dual β-lactam combinations exhibit synergism against *Mab* even in the absence of BLIs. For example, despite being rapidly hydrolyzed by the native β-lactamase Bla_Mab_, amoxicillin demonstrates strong synergism with β-lactams such as imipenem or ceftaroline. This suggests that the second β-lactam either acts as a functional BLI surrogate or targets complementary pathways. Supporting this, experiments using penicillin- and carbapenem-based probes revealed that β-lactams bind to multiple *Mab* proteins simultaneously, reinforcing the idea that their synergy arises from targeting complementary essential proteins. Finally, triple combinations comprising dual β-lactam and one BLI, such as amoxicillin + ceftaroline + avibactam, achieved very high synergy, underscoring the complementary roles of dual β-lactams and BLIs. The evidence in this study necessitates a revised model that can more accurately explain the activities of β-lactams and BLIs and underscores the potential for optimizing β-lactam/BLI regimens against *Mab*.

**IMPORTANCE:** This research challenges old assumptions about how antibiotics fight bacteria, particularly *Mycobacterium abscessus* (*Mab*), a tough-to-treat infection. Traditionally, β-lactam antibiotics were thought to stop bacterial growth, while β-lactamase inhibitors (BLIs) just protected them from breakdown. However, this study reveals that BLIs like clavulanic acid can work together with another BLI or β-lactam antibiotics for stronger effects. Surprisingly, even combinations comprising two BLIs can be highly effective, showing they target multiple critical bacterial processes simultaneously. Triple combinations—two β-lactams and one BLI—proved especially powerful. These findings overturn outdated ideas, offering a smarter way to use these drugs to combat difficult infections and save lives.

## INTRODUCTION

The peptidoglycan, a key structural component of the bacterial cell wall, is the target for more than 50% of antibiotic prescriptions today, specifically those in the β-lactam class (1). The traditional model of β-lactam action is largely based on studies of a few model organisms, including *E. coli, S. aureus, and B. subtilis*. According to this model, β-lactams exert their antibacterial effects by inhibiting enzymes essential for peptidoglycan synthesis, such as DD-transpeptidases (DDT, also known as penicillin-binding proteins or PBP) (2). These enzymes are required for bacterial cell survival, growth and division.

Bacteria commonly develop resistance to β-lactams through the production of β-lactamases, enzymes that hydrolyze and inactivate β-lactams (3). To counteract this, β-lactamase inhibitors (BLIs) are used in combination with β-lactams to treat infections caused by bacteria that produce β-lactamases. In this traditional model, BLIs are believed to only inhibit β-lactamases, thereby protecting the β-lactam, which retains its antibacterial activity in these combination treatments (4).

This model was challenged by evidence that demonstrated the presence of a new enzyme class that also catalyzes peptidoglycan synthesis but via a different mechanism than DDTs (5). While DDTs crosslink peptide side chains at 4→3 positions in the peptidoglycan, a distinct enzyme class forms crosslinks at 3→3 positions. This enzyme class, named LD-transpeptidase (LDT), was discovered in a clinical isolate of *Enterococcus faecium* resistant to ampicillin, a penicillin-class β-lactam (6). In this isolate, resistance was due to LDT activity rather than β-lactamases or mutations in DDTs. Further studies showed that while penicillin-class β-lactams are less effective against LDTs, carbapenems—a different subclass of β-lactams— are potent inhibitors of these enzymes (7–9).

The organism used in this study, *Mycobacterium abscessus* (*Mab*), employs both LDTs and DDTs for peptidoglycan synthesis with majority of the linkages generated by the LDTs (10). In a related and more commonly known organism, *Mycobacterium tuberculosis* (*Mtb*), LDTs also generate the majority of peptidoglycan crosslinkages (11–13). LDTs are not unique to mycobacteria; they are found in several clinically important bacteria, including *E. cloacae, E. coli, P. aeruginosa, K. pneumoniae,* and *C. difficile* (8, 14–17). Although 3→3 linkages in the peptidoglycan of *Mtb* and *E. coli* were first reported in 1974 and 1987, respectively (18, 19), the enzymatic basis for these linkages was overlooked, because the traditional model emphasized DDTs as the sole source of peptidoglycan crosslinking.

The presence of both LDTs and DDTs in *Mab*, along with their differing susceptibilities to various β-lactam subclasses, led to the hypothesis that combining two β-lactam antibiotics (also known as dual β-lactams) targeting both enzyme classes may produce synergistic antibacterial effects (8). Indeed, independent research has confirmed this hypothesis, showing mechanistic evidence of synergy between β-lactams with distinct chemical structures (8, 9, 20). This synergy demonstrates the presence of multiple non-redundant proteins that are bound by β-lactams.

The *Mab* genome encodes five LDTs, multiple DDTs, and several putative DDTs based on sequence homology (17, 21). Since the catalytic sites of LDTs and DDTs are chemically and structurally distinct, no single β-lactam may effectively inhibit all these enzymes (22). Additionally, the *Mab* genome encodes at least one β-lactamase, Bla_Mab_ (gene locus MAB_2875) (23, 24). In *Mtb*, in addition to its native β-lactamase, BlaC (gene locus Rv2068c), several DDTs also inactivate β-lactams (25), highlighting the ability of several peptidoglycan synthesis enzymes to inactivate β-lactams in addition to those enzymes that have been traditionally believed to be the only source of β-lactamase activity.

Based on the evidence that multiple LDTs, DDTs, and β-lactamases exist in *Mab,* in this study, we hypothesized that a combination comprising multiple β-lactams and BLIs would exhibit synergistic effects against *Mab*. The rationale behind this hypothesis is that the presence of multiple LDTs, DDTs, and β-lactamases necessitates the use of multiple β-lactams and BLIs to simultaneously inhibit different targets in peptidoglycan synthesis, thereby effectively blocking the entire pathway. This hypothesis challenges the traditional paradigm, which relies on combinations of only one β-lactam and one BLI in clinical treatments of *Mab* infection (24, 26). If proven correct, this approach could demonstrate untapped potential in using β-lactams and BLIs in novel combinations to treat *Mab* and other bacterial infections where multiple LDTs, DDTs, and β-lactamases are critical to disease progression. Therefore, the overall aim of this study was to not only assess if combinations of multiple β-lactams and BLIs are more potent than dual β-lactams but also to generate proof-of-concept that can support future studies to develop more potent combinatorial treatments with β-lactams and BLIs.

To test this hypothesis, we evaluated and compared the potencies of combinations of clinically relevant β-lactams from all three subclasses and BLIs from known classes against *Mab* using the checkerboard assay (27). This assay generates the fractional inhibitory concentration index (FICI) which represents the summation of the inhibition of microbial growth contributed by each drug in the combination. A stringent interpretation of the FICI was used, with FICI of ≤ 0.5 indicating synergy, >0.5 - <4 indicating indifference, and >4 indicating antagonism (28). A FICI of ≤ 0.5 results when two agents in the combination exist at only <0.25x their respective minimum inhibitory concentration (MIC) and yet inhibit growth of the microbe.

## RESULTS

### Imipenem exhibits synergy with most β-lactams in inhibiting *Mab* growth *in vitro*

To assess the activity of different dual β-lactam combinations, seven β-lactams representing three widely used subclasses were evaluated. These included amoxicillin and oxacillin from the penicillin subclass; cefoxitin, ceftazidime, and ceftaroline from the cephalosporin subclass; and doripenem and imipenem from the carbapenem subclass. The MIC of each β-lactam against *Mab* was determined to establish activity arising from each individual agent (**Table S1**). Whether 21 dual β-lactams that comprise all possible combinatorial pairs from the seven β-lactams exhibit synergism, indifference or antagonism in inhibiting *Mab* growth was assessed by determining their FICI. The FICIs for 15 dual β-lactams (71%) were ≤ 0.5, indicating synergistic effects in inhibiting *Mab* growth (**Figure 1a**). The mean FICI for combinations sharing one common β-lactam was calculated to identify the β-lactams that contributed most to synergism (**Figure 1b**). The dual β-lactams containing imipenem had the lowest mean FICI (0.26), demonstrating imipenem as the most common β-lactam in synergistic dual β-lactams. Therefore, the β-lactam that is most likely to exhibit synergy against *Mab* when combined with another β-lactam is imipenem. In contrast, dual β-lactams with oxacillin had the highest mean FICI (0.63), indicating that oxacillin exhibits synergy with the fewest β-lactams to inhibit *Mab* growth.

**Figure 1:**
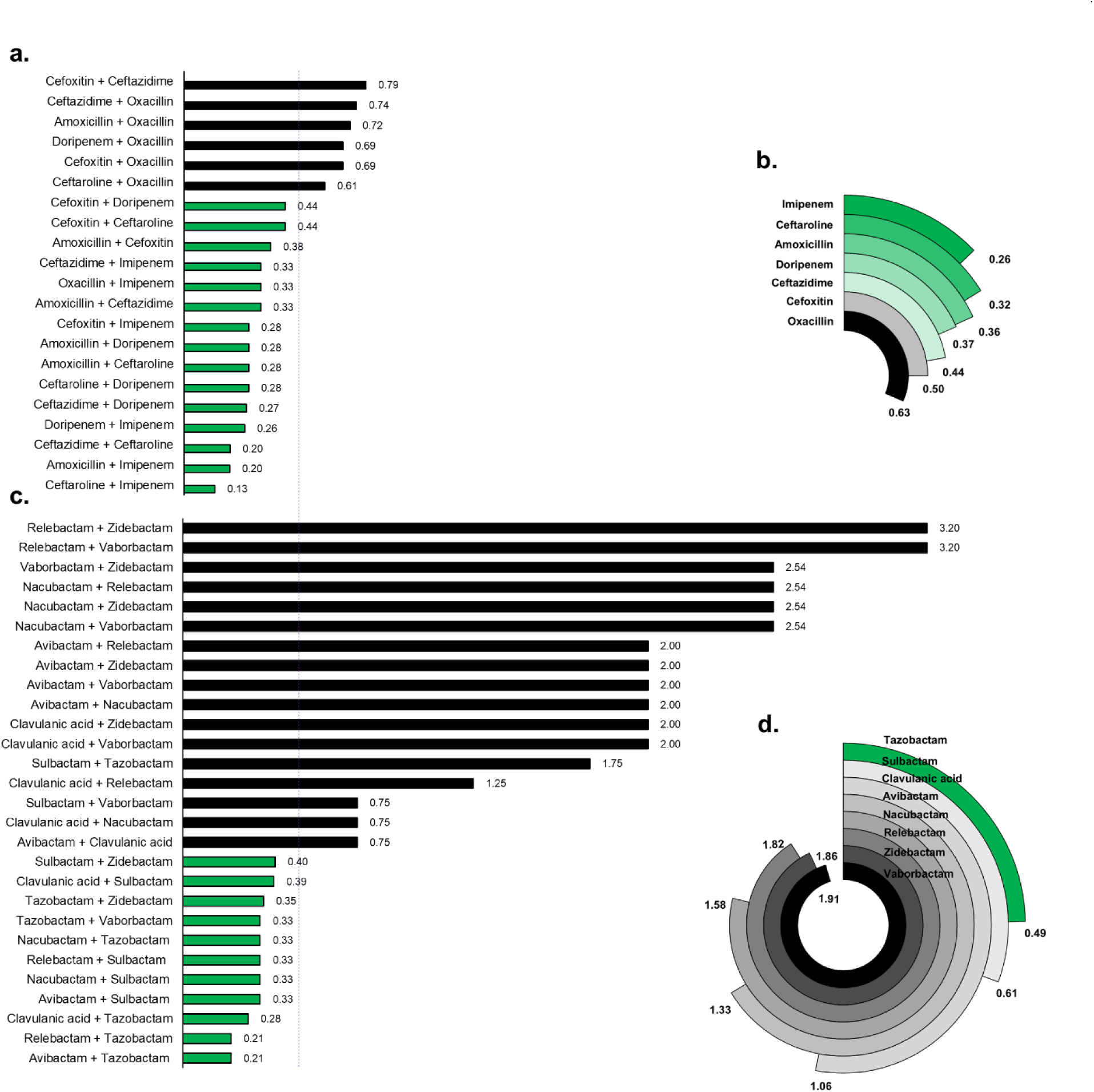
Activities of dual β-lactams or dual β-lactamase inhibitors (BLI) against *M. abscessus*. **a**) Fractional inhibitory concentration index (FICI) of 21 dual β-lactams arising from seven β-lactams are shown. A dotted line at FICI of 0.5 demarks combinations that exhibited synergy (shown in green) and those that lacked synergy (shown in black). 71% of dual β-lactams exhibited synergy. **b**) The mean FICIs for dual β-lactams sharing one common β-lactam, listed on the left side of this figure, are shown. **c**) FICI of 28 dual BLIs arising from eight BLIs are shown. 39% of dual BLIs exhibited synergy. **d**) The mean FICIs for dual BLIs sharing one common BLI are shown.

### Tazobactam synergizes with most β-lactamase inhibitors against *Mab* growth *in vitro*

Similarly, to determine the activities of dual BLI combinations (2BLI) against *Mab* growth, eight agents classified as BLIs were considered. Among them, clavulanic acid, sulbactam and tazobactam belong to the β-lactam chemical class. Avibactam, nacubactam, relebactam and zidebactam are non β-lactam bicyclic agents. Vaborbactam, also a non β-lactam, is a boronic acid agent (29). MICs of each BLI (**Table S1**) and FICIs of 28 dual BLIs arising from the BLI were determined against *Mab*. Eleven of these dual BLIs (39%) showed synergism, with FICI values ≤ 0.5 (**Figure 1c**). Surprisingly, one of the agents in all synergistic dual BLIs is a β-lactam-based BLI clavulanic acid, sulbactam or tazobactam. None of the dual BLIs comprised entirely of non-β-lactam BLI exhibited synergism. To identify the BLI that most frequently contributed to synergism, the mean FICI of pairs sharing one common BLI was calculated. Dual BLIs containing tazobactam had the lowest mean FICI (0.49), indicating its ability to exhibit synergy with the greatest number of BLIs in inhibiting *Mab* growth (**Figure 1d**). Conversely, pairs with vaborbactam had the highest mean FICI, highlighting its ability to exhibit synergism with a limited number of BLIs.

### Several β-lactam + β-lactamase inhibitor combinations exhibit synergism against *Mab* growth

To identify β-lactam and BLI combinations that exhibit synergism in inhibiting *Mab* growth, the FICIs of 56 unique combinations comprising one β-lactam and one BLI (1B+1BLI)—derived from the same seven β-lactams and eight BLIs considered above—were determined. Among these, 38 pairs (68%) had a FICI ≤ 0.5, indicating a synergistic effect in inhibiting *Mab* growth (**Figure 2a**). To pinpoint which β-lactams were most synergistic across different BLIs, the mean FICI of each β-lactam paired with all BLIs was calculated. Amoxicillin had the lowest mean FICI (0.09) and exhibited synergism with all BLIs (**Figure 2b**). Ceftaroline, imipenem, and doripenem also had mean FICIs below 0.5, indicating synergy with several BLIs. In contrast, ceftazidime, oxacillin, and cefoxitin showed synergy with the fewest BLIs, with ceftazidime being synergistic with only two. In summary, the β-lactams ranked by their synergy with BLIs in inhibiting *Mab* growth *in vitro* as follows: amoxicillin > ceftaroline > imipenem > doripenem > cefoxitin > oxacillin > ceftazidime.

**Figure 2:**
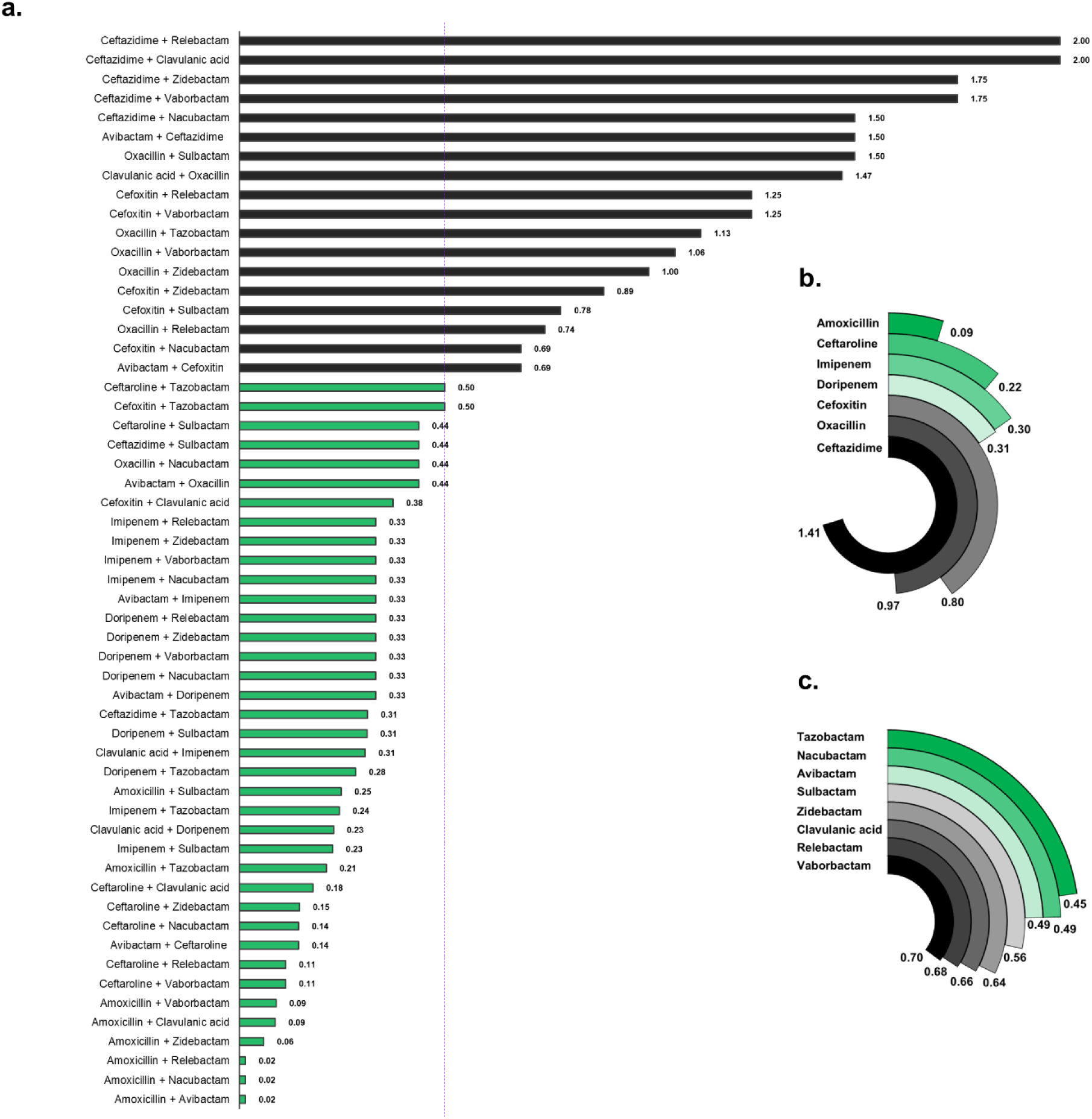
Activities of combinations comprising one β-lactam and one β-lactamase inhibitor (1B+1BLI) against *M. abscessus*. **a**) Fractional inhibitory concentration index (FICI) of 56 1B+1BLI arising from seven β-lactams and eight BLIs are shown. A dotted line at FICI of 0.5 demarks combinations that exhibited synergism (shown in green) and those that lacked synergism (shown in black). 68% of 1B+1BLI combinations exhibited synergy. **b**) The mean FICIs for 1B+1BLI combinations sharing one common β-lactam, listed on the left side of this figure, are shown. **c**) The mean FICIs for 1B+1BLI combinations sharing one common BLI are shown.

To identify the BLIs that exhibit synergism with most β-lactams, the mean FICIs of each BLI paired with various β-lactams were calculated (**Figure 2c**). Combinations containing tazobactam, nacubactam, or avibactam had mean FICIs < 0.5, indicating that these BLIs synergize with a broader range of β-lactams compared to the other BLIs. However, the differences in the number of β-lactams with which each BLI exhibited synergy were not sufficiently distinct to inform prioritization of one BLI over another.

### β-lactamase inhibitors exhibit synergism with dual β-lactams against *Mab*

We hypothesized that the anti-*Mab* activity of synergistic dual β-lactams could be further enhanced by the addition of a BLI. This hypothesis is based on the presence of multiple proteins encoded by *Mab* that serve as targets for both β-lactams and BLIs (7, 9, 10). To test this hypothesis, we evaluated *Mab* growth in the presence of synergistic β-lactam pairs combined with a BLI and calculated the FICIs of the combinations.

Among the 15 synergistic dual β-lactams identified above and eight BLIs, all 120 unique combinations of dual β-lactams + one BLI (2B+1BLI) were assessed. A checkerboard assay was conducted, with the dual β-lactams on one axis and a BLI on the other. Out of the 120 combinations tested, 110 (92%) showed FICI values ≤ 0.5, indicating that BLIs can indeed enhance the effectiveness of the majority of dual β-lactams in inhibiting *Mab* growth (**Table 1**).

**Table 1:** Activities of combinations comprising a dual β-lactam and a β-lactamase inhibitor against *M. abscessus* ATCC 19977. The fractional inhibitory concentration indexes (FICI) of each dual β-lactam + β-lactamase inhibitor combination are shown. FICIs ≤ 0.5 indicating synergism are highlighted in green. The last column lists the mean FICI for each dual β-lactam when combined with a β-lactamase inhibitor.

| Dual β-lactam | Sulbactam | Clavulanic acid | Tazobactam | Avibactam | Nacubactam | Vaborbactam | Zidebactam | Relebactam | Mean FICI |
| --- | --- | --- | --- | --- | --- | --- | --- | --- | --- |
| Amoxicillin + Ceftaroline | 0.186 | 0.069 | 0.143 | 0.016 | 0.023 | 0.050 | 0.075 | 0.023 | 0.073 |
| Amoxicillin + Imipenem | 0.171 | 0.084 | 0.126 | 0.016 | 0.098 | 0.035 | 0.098 | 0.098 | 0.091 |
| Amoxicillin + Ceftazidime | 0.186 | 0.148 | 0.186 | 0.098 | 0.098 | 0.100 | 0.140 | 0.098 | 0.132 |
| Ceftaroline + Doripenem | 0.133 | 0.122 | 0.143 | 0.145 | 0.145 | 0.145 | 0.145 | 0.145 | 0.140 |
| Amoxicillin + Doripenem | 0.290 | 0.102 | 0.264 | 0.098 | 0.098 | 0.094 | 0.132 | 0.091 | 0.146 |
| Cefoxitin + Ceftaroline | 0.232 | 0.143 | 0.132 | 0.145 | 0.145 | 0.145 | 0.155 | 0.145 | 0.155 |
| Ceftazidime + Ceftaroline | 0.184 | 0.167 | 0.176 | 0.145 | 0.145 | 0.145 | 0.165 | 0.145 | 0.159 |
| Amoxicillin + Cefoxitin | 0.347 | 0.176 | 0.259 | 0.098 | 0.098 | 0.100 | 0.129 | 0.098 | 0.163 |
| Imipenem + Oxacillin | 0.284 | 0.217 | 0.171 | 0.207 | 0.207 | 0.207 | 0.332 | 0.207 | 0.229 |
| Ceftaroline + Imipenem | 1.037 | 0.223 | 0.297 | 0.145 | 0.145 | 0.207 | 0.207 | 0.155 | 0.302 |
| Cefoxitin + Doripenem | 0.367 | 0.280 | 0.311 | 0.312 | 0.332 | 0.332 | 0.332 | 0.332 | 0.325 |
| Ceftazidime + Doripenem | 0.417 | 0.273 | 0.327 | 0.338 | 0.332 | 0.332 | 0.332 | 0.332 | 0.335 |
| Doripenem + Imipenem | 0.311 | 0.262 | 0.311 | 0.332 | 0.332 | 0.332 | 0.500 | 0.332 | 0.339 |
| Cefoxitin + Imipenem | 0.332 | 0.218 | 0.332 | 0.332 | 0.749 | 0.749 | 0.749 | 0.749 | 0.526 |
| Ceftazidime + Imipenem | 0.354 | 0.313 | 0.438 | 1.325 | 1.500 | 1.583 | 1.187 | 1.469 | 1.021 |

Among these, the dual β-lactams (amoxicillin+ceftaroline) and (amoxicillin+imipenem) showed the highest synergy when combined with avibactam, achieving FICI values of 0.016 for each combination. We calculated the mean FICIs for each dual β-lactam to identify those that synergized most frequently with different BLIs (**Table 1**). Based on the mean FICI values, the dual β-lactams ranked by their synergistic potential when combined with a BLI as follows: (amoxicillin+ceftaroline) > (amoxicillin+imipenem) > (amoxicillin+ceftazidime) > (ceftaroline+doripenem) > (amoxicillin+doripenem) > (cefoxitin+ceftaroline) > (ceftazidime+ceftaroline) > (amoxicillin+cefoxitin) > (imipenem+oxacillin) > (ceftaroline+imipenem) > (cefoxitin+doripenem) > (ceftazidime+doripenem) > (doripenem+imipenem) > (cefoxitin+imipenem) > (ceftazidime+imipenem). Notably, dual β-lactams containing amoxicillin were most frequently synergistic when paired with a BLI.

Among the 10 combinations that did not exhibit synergy, 4 (40%) were combinations of BLIs with (imipenem+cefoxitin), and 5 (50%) were combinations with (imipemen+ceftazidime). The addition of several BLIs to the dual β-lactams (imipenem+cefoxitin) and (imipemen+ceftazidime) failed to lower their FICI perhaps due to the high level of synergy between imipenem and cefoxitin (FICI = 0.28) and imipenem and ceftazidime (FICI= 0.33) (**Figure 1a**). This suggests that adding a BLI provides only marginal enhancement to the activities of these particular dual β-lactams.

### Adding a second β-lactamase inhibitor improves potencies of select β-lactam + β-lactamase inhibitor pairs

We evaluated whether the anti-*Mab* activity of synergistic dual BLIs could be enhanced by adding a β-lactam. Specifically, we tested *Mab* growth *in vitro* in the presence of combinations comprising one β-lactam and dual BLIs (1B+2BLI). A total of 77 unique 1B+2BLI combinations, derived from seven β-lactams and 11 synergistic dual BLI identified above, were assessed. The FICI indicated synergy (FICI ≤ 0.5) in 60 (78%) of these combinations (**Table 2**).

**Table 2:** Activities of combinations comprising dual β-lactamase inhibitors and a β-lactam against *M. abscessus* ATCC 19977. The fractional inhibitory concentration indexes (FICI) of each dual β-lactamase inhibitor + β-lactam combination are shown. FICIs ≤ 0.5 indicating synergism are highlighted in green. The last column lists the mean FICI for each dual β-lactamase inhibitor when combined with a β-lactam.

| Dual β-lactamase inhibitor | Amoxicillin | Cefoxitin | Ceftazidime | Ceftaroline | Doripenem | Imipenem | Oxacillin | Mean FICI |
| --- | --- | --- | --- | --- | --- | --- | --- | --- |
| Tazobactam + Avibactam | 0.016 | 0.217 | 0.332 | 0.107 | 0.163 | 0.749 | 0.162 | 0.249 |
| Clavulanic acid + Tazobactam | 0.143 | 0.311 | 0.290 | 0.176 | 0.176 | 0.332 | 0.332 | 0.252 |
| Tazobactam + Nacubactam | 0.031 | 0.280 | 0.332 | 0.072 | 0.186 | 0.749 | 0.228 | 0.268 |
| Tazobactam + Vaborbactam | 0.037 | 0.311 | 0.332 | 0.112 | 0.150 | 0.749 | 0.249 | 0.277 |
| Sulbactam + Avibactam | 0.016 | 0.263 | 0.749 | 0.104 | 0.193 | 0.749 | 0.228 | 0.329 |
| Sulbactam + Relebactam | 0.026 | 0.258 | 0.749 | 0.107 | 0.167 | 0.749 | 0.270 | 0.332 |
| Sulbactam + Nacubactam | 0.031 | 0.258 | 0.367 | 0.104 | 0.227 | 1.249 | 0.367 | 0.372 |
| Tazobactam + Zidebactam | 0.086 | 0.354 | 0.749 | 0.143 | 0.186 | 0.749 | 0.347 | 0.373 |
| Clavulanic acid + Sulbactam | 0.159 | 0.354 | 0.290 | 0.176 | 0.232 | 0.687 | 0.749 | 0.378 |
| Sulbactam + Zidebactam | 0.125 | 0.438 | 0.749 | 0.143 | 0.181 | 0.749 | 0.396 | 0.397 |
| Tazobactam + Relebactam | 1.070 | 0.280 | 0.332 | 0.072 | 0.186 | 0.749 | 1.070 | 0.537 |

This synergy rate marks a 10% increase over the 68% observed with 1B+1BLI combinations, suggesting that many 1B+1BLI pairs gain additional activity when a second BLI is added. For all β-lactams except imipenem, the addition of a second BLI almost always led to a synergistic 1B+2BLI combination. Amoxicillin demonstrated strong synergy with dual BLIs (lowest average FICI), with the exception of the relebactam+tazobactam pair. Cefoxitin, ceftaroline, and doripenem consistently exhibited synergy with all tested dual BLI combinations. Conversely, imipenem generally showed no additional activity (FICI > 0.5) when a second BLI was added, except in the combination clavulanic acid+tazobactam.

### Triple combinations comprising dual β-lactams + one BLI exhibit added synergy against *Mab*

As described above, the addition of a BLI to dual β-lactams or to combinations of 1β-lactam+1BLI leads to synergy in inhibiting *Mab* growth *in vitro*. This suggests that these triple combinations inhibit complementary targets in *Mab*. However, experiments described so far have not tested if these triple combinations saturate the available β-lactam and BLI targets in *Mab*.

To explore this idea, we hypothesized that combining four agents—two β-lactams and two BLIs (2B+2BLI)—may saturate available targets and manifest as enhanced synergism. To test this hypothesis, we evaluated the activities of 165 unique 2B+2BLI combinations derived from 15 synergistic dual β-lactams and 11 synergistic dual BLIs using the checkerboard assay. For 2B+2BLI to achieve synergy, each component would have to be present at ≤1/16x their respective MICs for the net FICI to be ≤ 0.5. Of the 165 2B+2BLI combinations, 130 (79%) were not synergistic. The FICI was ≤ 0.5 for 35, all of which included amoxicillin as the common β-lactam (**Table S2**). Therefore, except for 2B+2BLI combinations that included amoxicillin, it appears that combinations comprising three agents, whether dual β-lactam + β-lactamase inhibitor, or a β-lactam + dual β-lactamase inhibitor may be sufficient to saturate available targets in *Mab*.

### β-lactam probes bind to multiple proteins in *Mab*

The findings above suggest that β-lactams and BLIs inhibit complementary targets in *Mab* to achieve synergy. However, the assays based on assessing *Mab* growth inhibition in the presence of β-lactams and BLIs do not directly confirm the binding of these agents to multiple targets in *Mab*. To generate direct evidence of proteins in *Mab* that are bound by β-lactams, as a proof-of-concept, we labeled *Mab* lysates with a recently described red fluorescent meropenem-Cy5, a β-lactam of the carbapenem subclass (30). To represent the penicillin subclass, we used the commercially available green fluorescent Bocillin-FL probe. These probes were incubated with *Mab* whole-cell lysate, and the resulting mixtures were analyzed using SDS-PAGE (**Figure 3**). Fluorescence scans of the gels reveal multiple bands, each corresponding to a unique protein bound by the probes. The meropenem probe revealed many enzymes not labeled by Bocillin-FL. While several proteins were bound by probes, variable band intensities suggest differential affinity of the β-lactam probes to the target proteins. These results provide direct evidence that there are many targets of β-lactams in *Mab* and that different β-lactam subclasses bind to multiple proteins in *Mab* but with distinct affinities and selectivities.

**Figure 3.**
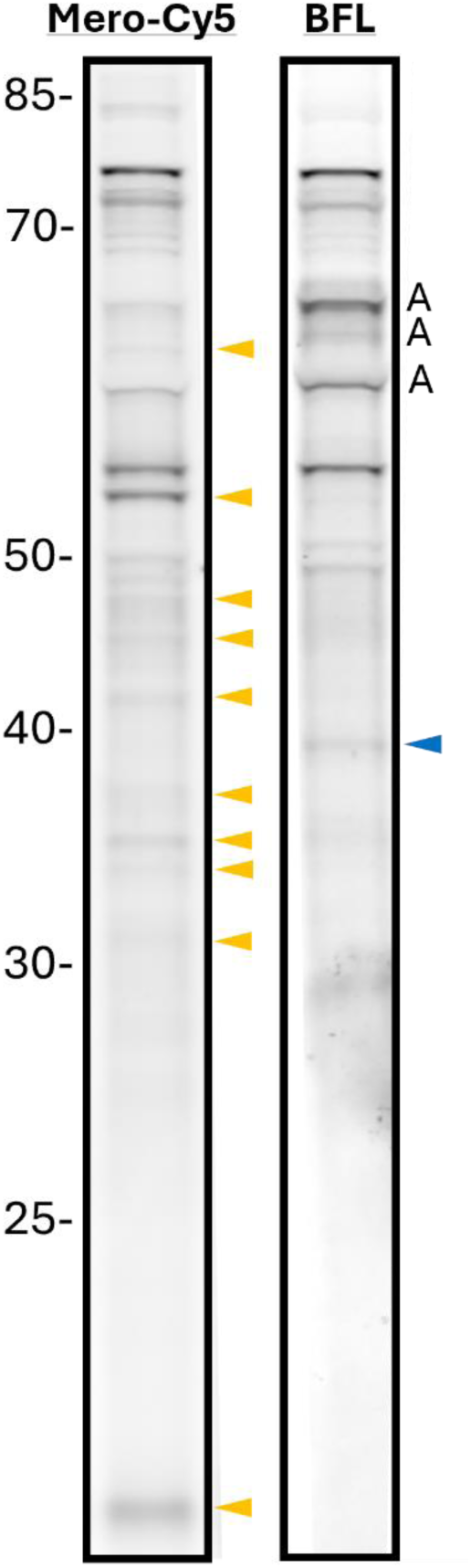
β-lactam probes bind multiple proteins in *Mab* lysates. *Mab* whole-cell lysate was treated with meropenem-Cy5 (Mero-Cy5) or Bocillin-FL (BFL), resolved using SDS-PAGE, and imaged. Three auto fluorescent bands (A) were observed in the green channel (BFL) but none were observed in the Cy5 channel. Targets uniquely identified by Mero-Cy5 are marked with a yellow arrow, while those identified by BFL are marked with a blue arrow. The markers on the left denote molecular weight of the proteins in kilo daltons.

## DISCUSSION

The historical model describing the mechanistic basis of β-lactams and BLIs posits that β-lactams inhibit microbial growth, while BLIs serve solely to protect β-lactams from degradation by β-lactamases (4). This framework has guided the development of combinations of one β-lactam and one BLI for treating bacterial infections. However, evidence from this study demonstrates that this model does not fully apply to *Mab*. Notably, numerous instances of synergism were observed against *Mab* in combinations that deviate from the traditional one β-lactam–one BLI paradigm.

The first line of evidence is the synergistic activity of select dual β-lactams against *Mab*, even in the absence of a BLI (**Figure 1a**). The relevance of β-lactamase activity to this synergy remains unclear. For example, amoxicillin is inactivated in the presence of the β-lactamase Bla_Mab_ (31), which explains its high MIC against *Mab* (**Table S1**). Despite this, amoxicillin exhibits strong synergism when paired with β-lactams such as imipenem, doripenem, cefoxitin, ceftazidime, or ceftaroline (**Figure 1a**). This raises the question of whether the second β-lactam in these pairs acts as a functional BLI for amoxicillin. Notably, the multiple protein targets bound by meropenem-Cy5 and bocillin-FL probes (**Figure 3**) may hold the answer to this conundrum-it suggests that synergism of dual β-lactams is achieved by simultaneous inhibition of several complementary targets.

The second line of evidence challenges the traditional role of BLIs as β-lactamase inhibitors without intrinsic antibacterial activity. Synergistic dual BLIs were identified, particularly those involving β-lactam-based agents such as clavulanic acid, sulbactam, and tazobactam (**Figure 1c**). These findings suggest that these agents may also inhibit essential targets in *Mab* beyond β-lactamase activity. For instance, in a prior study to assess β-lactam targets in *M. tuberculosis*, pre-treatment with clavulanic acid blocked labeling of its β-lactamase, BlaC, by meropenem-Cy5. However, labeling of several other proteins was also lost, suggesting that clavulanic acid inhibits multiple targets (30). Historically regarded as distinct from β-lactams, these BLIs may share antibacterial properties due to their chemical composition as β-lactams. For example, the sulbactam+durlobactam combination is an example of a dual BLI, recently approved by the FDA for bacterial pneumoniae, which likely inhibits proteins essential for cell survival in addition to inhibiting β-lactamases (32).

Tazobactam, frequently found in synergistic combinations (mean FICI = 0.49), appears to have a broader spectrum of activity against *Mab* targets compared to other BLIs. Previous studies indicate that tazobactam, sulbactam, and clavulanic acid do not inhibit Bla_Mab_ (23), implying the presence of additional proteins targeted by tazobactam. As tazobactam exhibited synergy with several β-lactams in our study, this evidence indicates the presence of β-lactamase activity additional to Bla_Mab_ that is relevant to *Mab* growth and is inhibited by tazobactam. Conversely, vaborbactam demonstrated limited synergy, likely reflecting its limited specificity for *Mab*’s β-lactamase repertoire.

Adding a BLI to synergistic dual β-lactams provided variable enhancement, with the greatest synergy observed in amoxicillin-based combinations. For instance, imipenem-cefoxitin showed strong baseline synergy (FICI = 0.28), leaving limited room for improvement. A prior study that assessed synergism of several dual β-lactams against 21 *Mab* clinical isolates found imipenem+cefoxitin exhibited synergy against 100% of the isolates (33). In contrast, amoxicillin combinations exhibited greater enhancement, likely due to BLIs restoring activity by neutralizing enzymatic degradation. Notably, triple combinations such as amoxicillin+ceftaroline+avibactam achieved exceptional synergy (FICI = 0.016), underscoring the complementary roles of β-lactams and BLIs.

Synergism was also observed in combinations involving 1B+2BLI, with 60 of 77 (78%) combinations achieving FICI ≤ 0.5—a 10% increase over 1B+1BLI combinations. For example, amoxicillin+clavulanic acid+nacubactam exhibited synergy, likely due to complementary inhibition of multiple β-lactamase classes. In contrast, among 165 2B+2BLI combinations, only those that contained amoxicillin produced synergy. This finding suggests redundancy of the fourth agent except for combinations containing amoxicillin.

Based on the emerging evidence from this study and prior research (8, 9, 26, 34–40), we propose a revised model to more accurately describe the activities of β-lactams and BLIs in *Mab* (**Figure 4**). In this model, each β-lactam targets multiple proteins critical for peptidoglycan synthesis, with each β-lactam in a synergistic pair generating complementary inhibition required to achieve optimal inhibition of essential targets. Furthermore, agents historically classified as BLIs—particularly those with β-lactam-based structures such as clavulanic acid, sulbactam, and tazobactam—are likely not restricted to β-lactamase inhibition alone. Instead, our findings suggest that synergistic dual BLIs also target essential components of peptidoglycan synthesis, contributing directly to bacterial growth inhibition.

**Figure 4.**
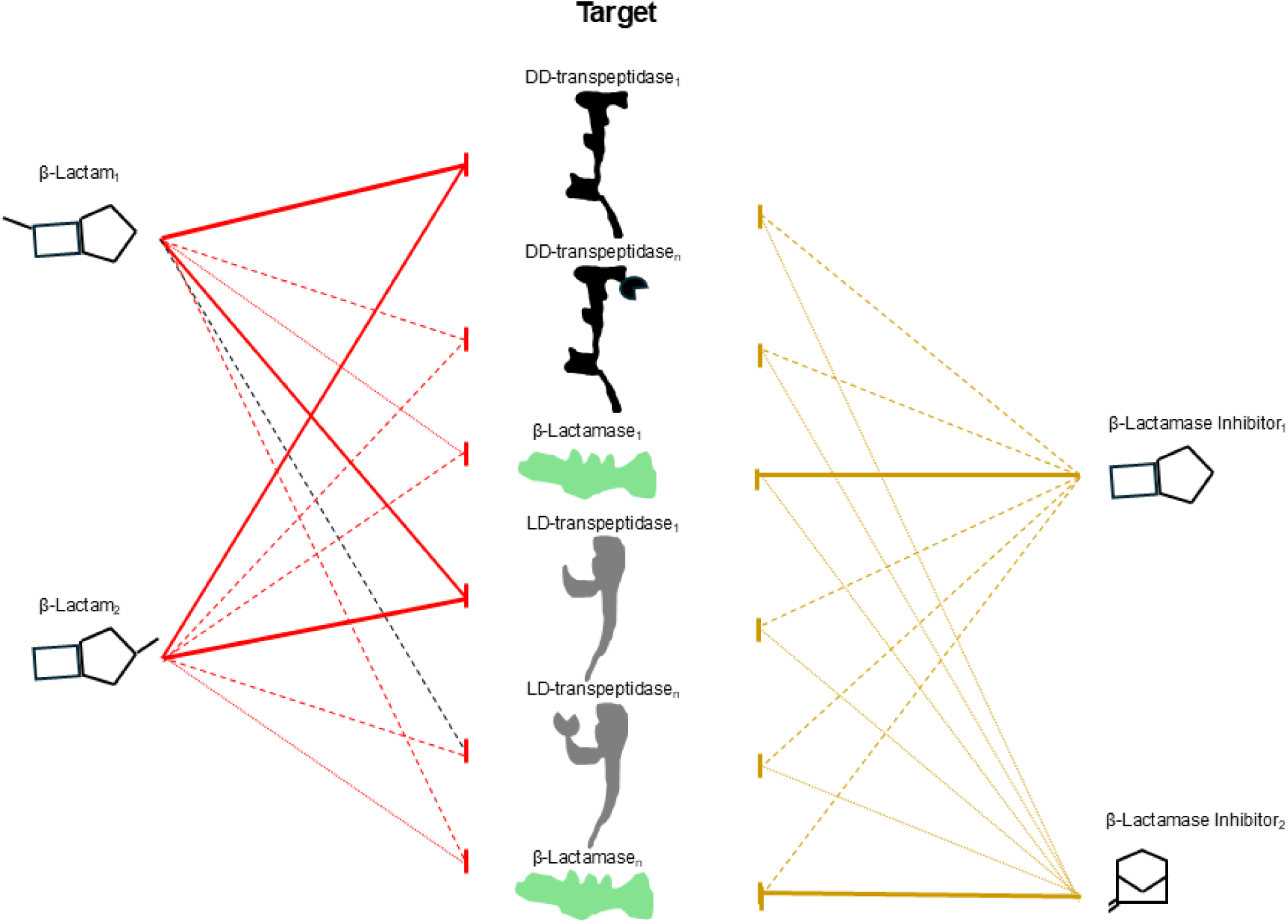
This model outlines the roles of β-lactams and β-lactamase inhibitors in targeting enzymes in *M. abscessus*. β-lactam_1_ and β-lactam_2_ represent two distinct β-lactams. They bind to an overlapping set of targets but exhibit varying efficiencies in inhibiting specific targets. Similarly, β-lactamase inhibitor_1_ and β-lactamase inhibitor_2_ represent two distinct β-lactamase inhibitors with each inhibiting overlapping targets but with varying efficiency. β-lactamase inhibitor_1_ is based on a β-lactam chemical structure, whereas β-lactamase inhibitor_2_ represents non β-lactam-based structure. Target enzymes: DD-transpeptidase_1_ and DD-transpeptidase_n_ depict the first and the n^th^ DD-transpeptidase. Similarly, LD-transpeptidase_1_ and LD-transpeptidase_n_ depict the first and the n^th^ LD-transpeptidase, and β-lactamase_1_ and β-lactamase_n_ depict the first and the n^th^ β-lactamase in *M. abscessus.* The thickness of the lines connecting β-lactams and β-lactamase inhibitors to their targets indicates the strength of inhibition.

In conclusion, the evidence from this study challenges the simplistic view that β-lactams target PBPs and LDTs essential for peptidoglycan synthesis, while BLIs merely inhibit β-lactamases. Instead, both β-lactams and BLIs appear to inhibit complementary targets in *Mab*. These findings highlight the potential to optimize β-lactam/BLI regimens, including novel triple combinations, to treat *Mab* infections. Future research should explore these mechanisms further, paving the way for more effective therapeutic strategies.

## MATERIALS AND METHODS

### Bacterial strains, drugs, culture media and *in vitro* growth conditions

*Mycobacterium abscessus (Mab)* strain ATCC 19977, isolated in 1950, commonly included as a reference for *Mab* in laboratory settings, was used in this study (41). The strain was procured from the American Type Culture Collection (Manassas, VA) and authenticated by genome sequencing (33). It was cultured in Middlebrook 7H9 broth (Difco, catalog no. 271310) supplemented with 0.5% glycerol, 10% albumin-dextrose-salt enrichment, and 0.05% Tween-80, at 37 °C in an orbital shaker at 220 RPM as described (42). However, for all assays involving determination of drug activities, Middlebrook 7H9 broth supplemented with 0.5% glycerol, 10% albumin-dextrose-salt enrichment without Tween-80 was used in U-bottom 96-well plates with 350 μL capacity per well. Powdered form of drugs were purchased from commercial vendors as follows: amoxicillin (Sigma-Aldrich, catalog no. A8523), oxacillin (Sigma-Aldrich, catalog no. 28221), cefoxitin (Sigma-Aldrich, catalog no. C4786), ceftazidime (Sigma-Aldrich, catalog no. C3809), ceftaroline fosamil hydrate (Sigma-Aldrich, catalog no. SML3102), imipenem (Sigma-Aldrich, catalog no. I0160), doripenem (Sigma-Aldrich, catalog no. SML1220), sulbactam (Sigma-Aldrich, catalog no. S9701), tazobactam (Sigma-Aldrich, catalog no. T2820), clavulanic acid (BOC Sciences, catalog no. B0084-073058), avibactam (Sigma-Aldrich, catalog no. SBR00075), relebactam (MedChemExpress, catalog no. HY-16752), vaborbactam (MedChemExpress, catalog no. HY-19930), zidebactam (MedChemExpress, catalog no. 120859), and nacubactam (MedChemExpress, catalog no. 109008).

### Determination of Minimum Inhibitory Concentration (MIC) of a drug

The MICs of β-lactams, amoxicillin, oxacillin, cefoxitin, ceftazidime, ceftaroline fosamil hydrate, imipenem, and doripenem, and β-lactamase inhibitors, sulbactam, tazobactam, clavulanic acid, avibactam, relebactam, vaborbactam, zidebactam, and nacubactam, against *Mab* were determined using a broth microdilution assay in accordance with the Clinical and Laboratory Standards Institute (CLSI) guidelines (43). A 10 mg/mL stock solution of each drug was made by dissolving in dimethyl sulfoxide (DMSO), which was used to make working solutions at lower dilutions in Middlebrook 7H9 broth supplemented with 0.5% glycerol, 10% albumin-dextrose-salt enrichment without Tween-80. Working drug solutions were diluted 2-fold serially in Middlebrook 7H9 broth to generate concentrations ranging from 512 μg/mL to 0.06 μg/mL. Using *Mab* grown to exponential phase, 10^5^ CFU of *Mab* was inoculated into each well in 200 μL final volume per well of 96-well culture plate. Two wells containing Middlebrook 7H9 broth only and two wells containing broth inoculated with 10^5^ CFU of *Mab* were included as negative and positive controls, respectively, and incubated at 30 °C for 72 h without shaking in accordance to the CLSI guidelines (43). Growth or lack thereof of *Mab* was assessed using Sensititre Manual Viewbox. The lowest concentration of the drug at which well suspension appeared identical to the well with broth only was recorded as the MIC of the drug. Two biological replicates with freshly grown *Mab* cultures and two technical replicates within each assay were performed. The final MIC was calculated as the mean of the MICs in biological and technical replicates.

### Determination of Fractional Inhibitory Concentration (FICI)

The checkerboard assay was used to determine the activities of combinations of drugs. This assay is a modification of the standard broth microdilution assay used for MIC determination, and was performed as described (27). The most stringent guideline for interpreting FICI was used: an FICI of ≤ 0.5 was interpreted as synergy, > 0.5 to 4 was interpreted as indifference, and > 4 as antagonism (28).

### Determination of FICI for two drugs (B+B or BLI+BLI or B+BLI)

To determine the FICIs of two drugs, working solutions of two drugs were added to Middlebrook 7H9 broth, each starting at 4-fold MIC and serially diluted up to 1/64-fold MIC allowing for all possible 2-fold dilution combinations ranging from 4- to 1/64-fold MIC combinations of each drug. The following drugs classified as β-lactams - amoxicillin, oxacillin, cefoxitin, ceftazidime, ceftaroline fosamil hydrate, imipenem, and doripem, and β-lactamase inhibitors - sulbactam, tazobactam, clavulanic acid, avibactam, relebactam, vaborbactam, zidebactam, and nacubactam-were included. All 21 unique β-lactam+ β-lactam (B+B) combinations, 28 unique β-lactamase inhibitor+ β-lactamase inhibitor (BLI+BLI) combinations, and 56 unique β-lactam+ β-lactamase inhibitor (1B+1BLI) combinations were evaluated. Using *Mab* grown to exponential phase, 10^5^ CFU of *Mab* was inoculated into each well in 200 μL final volume per well of 96-well culture plate. Two wells containing Middlebrook 7H9 broth only and two wells containing broth inoculated with 10^5^ CFU of *Mab* were included as negative and positive controls, respectively, and incubated at 30 °C for 72 h without shaking in accordance to the CLSI guidelines (43). Growth or lack thereof of *Mab* was assessed using Sensititre Manual Viewbox. The FICI of drug combinations were calculated as described (27): FICI_XY_ = FICI_X_ + FICI_Y_= ([X] /MIC_X_) + ([Y] / MIC_Y_). MIC_X_ and MIC_Y_ are MICs of drugs X and Y, respectively, when used alone. [X] and [Y] are the concentrations of the drugs X and Y, respectively, in the first well in decreasing drug concentrations where *Mab* growth is not observed. As growth is not observed in multiple wells in two axes, mean FICI_XY_ is calculated from the FICI_XY_ from each well and reported as the final FICI_XY_.

### Determination of FICI for three drugs (2B+1BLI or 1B+2BLI) or four drugs (2B+2BLI)

Identical protocol as described above for two agents was used with the exception of combining two drugs in one axis. For instance, to determine FICIs of combinations comprising two β-lactams and one β-lactamase inhibitor (2B+1BLI), all 120 unique combinations arising from 15 dual β-lactams (2B) that exhibited synergism (**Figure 1a**) and all 8 β-lactamase inhibitors were considered. Dual β-lactams with each β-lactam at 4-fold its MIC were combined. This combination and β-lactamase inhibitor at 4-fold its MIC were serially diluted in the same way as described for two drug combinations. The FICI of three drug combinations were calculated as described (27): FICI_XYZ_ = FICI_X_ + FICI_Y_ + FICI_Z_ = ([X] /MIC_X_) + ([Y] / MIC_Y_) + ([Z] / MIC_Z_).

Similarly, to determine the FICI for four drugs comprising two β-lactams and two β-lactamase inhibitor (2B+2BLI), all 165 unique combinations arising from 15 dual β-lactams (2B) that exhibited synergism (**Figure 1a**) and 11 dual β-lactamase inhibitor that exhibited synergism (**Figure 1c**) were considered. Dual β-lactams with each β-lactam at 4-fold its MIC were combined. Similarly, Dual β-lactamase inhibitors with each β-lactamase inhibitor at 4-fold its MIC were combined. The dual β-lactam and dual β-lactamase inhibitor combinations, starting at 4-fold its MIC were serially diluted in the same way as described for two drug combinations. The FICI of four drug combinations were calculated as described (27): FICI_WZYZ_ = FICI_W_ + FICI_X_ + FICI_Y_ + FICI_Z_ = ([W] /MIC_W_) + ([X] /MIC_X_) + ([Y] / MIC_Y_) + ([Z] / MIC_Z_). For a four-drug combination to achieve synergy, each component would have to be present at ≤1/16x their respective MICs for the net FICI to be ≤0.5.

### Fluorescent labeling of β-lactam targets

Meropenem-Cy5 (Mero-Cy5) was generated through a Cu-catalyzed azide-alkyne click reaction as previously reported (30). Bocillin FL (Bo FL, Invitrogen, catalog no. 13233) was purchased commercially and handled according to manufacturer protocols. *Mab* 19977 whole-cell lysate (15 μg total protein) was labeled with 5 μM Mero-Cy5 or Bo FL (60 min, RT, in dark). Labeled lysates were resolved via SDS-PAGE, fixed, and imaged on a Typhoon multimodal imager (Cytiva).

## FUNDING

This study was supported by NIH awards R01 AI 155664 to GL and R01 AI 149737 to KEB.

## AUTHOR CONTRIBUTIONS

BR: methodology, study design, investigation, data analysis and interpretation, manuscript preparation. YX and CMP: methodology and investigation. KLD: methodology, study design, investigation, data analysis and interpretation, manuscript preparation. KEB: study design, project administration, data interpretation, manuscript preparation, and funding acquisition. GL: study conception, study design, project administration, data interpretation, manuscript preparation, and funding acquisition.

